# Bidirectional modulation of neuronal excitability via ionic actuation of potassium

**DOI:** 10.1101/2022.04.03.486896

**Authors:** Claudio Verardo, Renaud Jolivet, Leandro Julian Mele, Michele Giugliano, Pierpaolo Palestri

## Abstract

Potassium K^+^ is a fundamental actor in the shaping of action potentials, and its concentration in the extracellular microenvironment represents a crucial modulator of neural excitability. Yet, its employment as a neuromodulation modality is still in its infancy. Recent advances in the technology of ionic actuators are enabling the control of ionic concentrations at the spatiotemporal scales of micrometers and milliseconds, thereby holding the promise of making the control of K^+^ concentration a key enabling technology for the next generation of neural interfaces. In this regard, a theoretical framework to understand the possibilities and limits offered by such technology is pivotal. To this aim, we exploit the Hodgkin-Huxley modeling framework, augmented to account for the perturbation of extracellular K^+^ concentration. We leverage methods of bifurcation analysis to investigate which regimes of electrical activity arise in the space of the input variables, namely the extent of ionic actuation and the synaptic current. We show that, depending on the type of target neuron, switchings of the class of excitability may occur in such space. These effects could rule out the possibility of eliciting tonic spiking when the extracellular K^+^ concentration is assumed as a sole control input. Building upon these findings, we show in simulations how to address the problem of neuromodulation via ionic actuation in a principled fashion. In this respect, we account for a bidirectional scenario, namely from the perspective of both inhibiting and stimulating electrical activity. We then provide a first-order motivation for the switchings of neural excitability in terms of the conductances of the K^+^-selective channels. Finally, we introduce a Pinsky-Rinzel-like model to investigate the effects of performing the ionic actuation locally at the neural membrane.

**Author summary:** Neural interfaces rely on technologies to sense and perturb the electrical activity of neurons. For the latter aim, many strategies have been established to date, each one targeting a different actor involved in the electrophysiology of neurons. Examples include electrical, chemical, and optogenetic techniques. However, the main actors that shape neuronal signals, namely ions such as potassium K^+^, are still not directly targeted. Recent advances in bioelectronic technologies are enabling the manipulation of ionic concentrations as a viable strategy for neuromodulation, which we refer to as *ionic actuation*. These findings come mainly from the experimental literature, and the theoretical understanding of how ionic actuation can be used to shape neural activity is still lacking. This paper aims to help fill this gap, adopting the ionic actuation of K^+^ as a case study. Our results could guide the design and control of novel neural interfaces targeting the ionic composition of cellular fluids. Moreover, they may suggest novel therapies for pathologies related to impairments in the regulation of ionic homeostasis, such as drug-resistant epilepsy.

## Introduction

The electrical activity of neurons relies on the correct balance and movement of various ionic species. Indeed, it emerges from the interplay of the homeostatic regulation of their concentrations in the cellular fluids, as well as their electrodiffusive transport across the neural membrane [1]. When transmembrane fluxes balance each other, steady-state conditions are established. Otherwise, instability arises and generates action potentials (also denoted as spikes). Once understood, one may attempt to artificially induce the desired electrical activity via these underlying phenomena, either directly or indirectly, paving the way to *neuromodulation devices* [2]. We point out that the term neuromodulation is used in this context to refer to a generic strategy able to perturb the electrical excitability of neurons and not to the more specific action of neuromodulators (e.g., serotonin, acetylcholine, dopamine) in the nervous system.

The possible strategies to accomplish neuromodulation are manifold and reflect the plethora of physiological actors that determine neural activity. Chemical methods involve the release of neurotransmitters such as GABA or glutamate in the extracellular microenvironment [3]. Electrical methods rely on the injection of electric currents in the nervous tissue through electrodes [4]. Optogenetics can also be deployed to this aim, expressing light-sensitive proteins able to induce transmembrane ionic fluxes in neurons [5]. Notwithstanding, the central role played by ions in neural electrophysiology suggests the direct manipulation of the ionic milieu as a further neuromodulation strategy. Indeed, a change in the ionic concentrations unbalances the transmembrane drift and diffusion fluxes, possibly affecting the excitability of nerve cells. In what follows, we shall refer to this technique as *ionic actuation*.

Arguably, one of the foremost candidates for ionic actuation is potassium K^+^ [6]. Indeed, an increase (respectively, decrease) of K^+^ extracellular concentration from its physiological level induces a depolarization (respectively, hyperpolarization) of the membrane potential. As a consequence, K^+^ may act as a modulator of the activity of neurons and other brain cells. For instance, a large body of literature reports oscillations of K^+^ as one of the main causes of the firing patterns observed in epilepsy and spreading depression, both experimentally [7–10] and computationally [11–18]. Further, K^+^ is a driver of the motility of the brain’s immune cells (microglia) [19] and is involved in sleep-wake cycles [20]. Such findings have recently suggested K^+^ concentration as a possible control variable to steer neurons to desired regimes of electrical activity [21,22]. In this work, we aim to further explore this idea, considering an ionic actuation strategy where K^+^ is the ionic species under control, as depicted in Fig. 1 (left).

**Fig 1.**
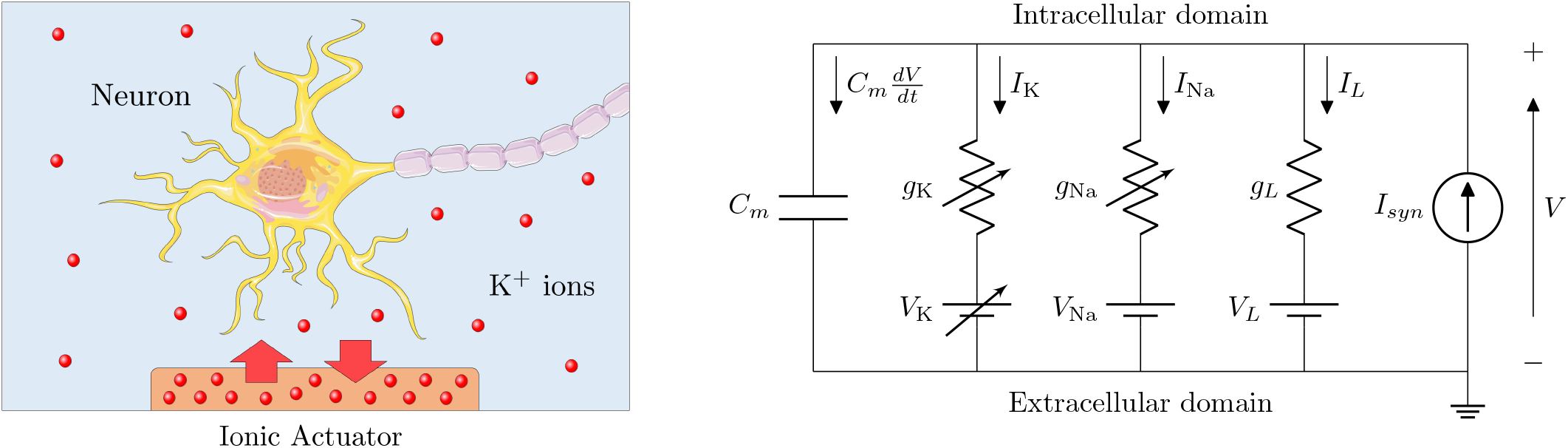
Neuromodulation via ionic actuation of potassium K^+^. Left) cartoon of the neuromodulation system under study. A K^+^-selective ionic actuator is placed in contact with the extracellular fluid nearby a neuron. It allows perturbing the local extracellular concentration of potassium, thereby steering the neuron to the desired regime of electrical activity. Right) equivalent Hodgkin-Huxley model employed in the analysis of the system. It is endowed with a variable K^+^ reversal potential *V*_K_ = *V*_K,0_ + Δ*V*_K_, see Eq. (2), to account for the release or uptake of ions in the extracellular milieu. Further, a parallel current *I_syn_* is added to explore the excitatory or inhibitory action of synapses (or other external stimuli), which are regarded as uncontrolled input of the system under study.

Recent technological advances gave rise to integrated devices able to deliver and uptake ionic species at spatiotemporal resolutions of micrometers and milliseconds, which we shall refer to as *ionic actuators*. They enable the implementation of the previously mentioned speculations in realistic applications of neural engineering. Examples of such devices have stemmed from iontronics [23] and have been successfully deployed to deliver K^+^ ions into nervous tissues, both *in-vitro* [24] and *in-vivo* [25]. These devices can also be deployed to deliver other charged molecules, such as the neurotransmitter GABA [26–31]. However, such neuromodulation strategies fundamentally differ from the aforementioned ionic actuation in the mechanism that affects the membrane potential. Indeed, neurotransmitters target specific binding sites to open ionic channels at synapses, while the manipulation of K^+^ concentration directly alters the drift-diffusive fluxes across the K^+^-selective channels.

To guide the design and control of ionic actuators in a neuromodulation scenario, a theoretical framework to assess the impact of ionic actuation on the electrical activity of neurons is pivotal. For instance, it would be helpful to identify which regimes of electrical activity can be reached by perturbing the K^+^ concentration. Moreover, it would be desirable to obtain a budget of the amount of K^+^ release or uptake required to eventually elicit such regimes in a neuron. Due to the high non-linearity of the neural membrane’s dynamics, the answers to such questions are not readily available. The present work aims to help bridge this gap, exploiting the Hodgkin–Huxley (HH) framework and methods of bifurcation analysis [32,33]. Together, they establish a principled framework to guide the inhibition and stimulation of electrical activity via the proposed neuromodulation modality.

The paper is structured as follows. First, we introduce the single-compartment model deployed in our main investigations. It is an HH-type model which is augmented to account for the presence of ionic actuation and synaptic activity, namely the input variables of the system. We then leverage bifurcation analysis to find which regimes of electrical activity arise in the space of the input variables. This analysis reveals possible switchings of the class of neural excitability that may rule out the triggering of tonic spiking via ionic actuation. Building upon these findings, we show how the system can be ionically steered to the desired regime in a principled fashion, regardless of the presence of a synaptic current. We further propose an interpretation of the switchings of the class of excitability based on the effect of the conductances of the K^+^-selective channels. Finally, we investigate the possibility of a spatially localized ionic actuation deploying a Pinsky-Rinzel-like two-compartment model.

## Methods

To investigate the effects of the proposed neuromodulation via ionic actuation of potassium, we employ the Hodgkin-Huxley (HH) formalism [34, 35] to model the electrical excitability of a neuron, as depicted in Figure 1 (right). For the sake of simplicity, we consider only K^+^ and Na^+^ channels responsible for the generation of action potentials. We denote with [K^+^]_*o*_ and [K^+^]_*i*_ the extracellular and intracellular concentrations of K^+^ in physiological conditions, i.e., in absence of ionic actuation. Instead, Δ[K^+^]_*o*_ is the perturbation of the extracellular concentration of K^+^ induced by the ionic actuator, which is assumed as a controlled input to the model. The effect of Δ[K^+^]_*o*_ is lumped in the model through a variable K^+^ reversal potential *V*_K_, which can be recasted according to the Nernst equation [1] as:

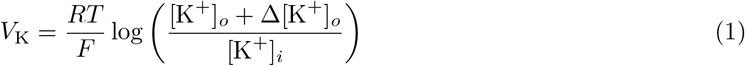

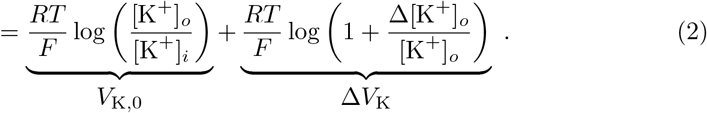

Here, *T* is the temperature, *R* the gas constant, and *F* the Faraday constant. The term *V*_K,0_ is the reversal potential under physiological conditions, while Δ*V*_K_ represents its perturbation during the action of the ionic actuator, and depends logarithmically on the fractional variation of extracellular concentration of K^+^. Being that the HH equations are linear in *V*_K_, we use Δ*V*_K_ in place of Δ[K^+^]_*o*_ as control variable in the forthcoming derivations, keeping in mind that we can map the former into the latter according to

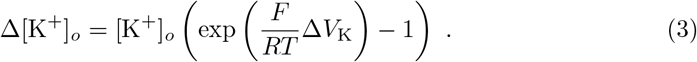

We further augment the HH model with an inward current *I_syn_* to investigate the effect of synaptic activity on the electrical excitability of the neuron. An excitatory synaptic current *I_syn_* > 0 depolarizes the membrane potential *V* and is expected to facilitate the generation of action potentials via Δ*V*_K_. The converse applies to an inhibitory synaptic current *I_syn_* < 0. Even though we refer to *I_syn_* as synaptic current, we point out that it may equivalently represent other external stimuli.

We assume first-order kinetics for the gating variables *x* ∈ {*n, m, h*}, which are ruled by the (in)activation curves *x*_∞_(*V*) and the time constants *τ_x_*(*V*). Along with the current balance equation, they provide the full set of equations of the system:

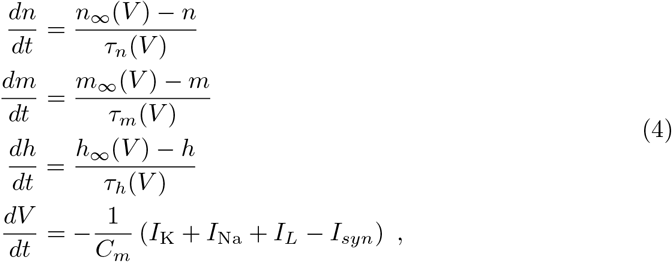

where *C_m_* is the membrane capacitance. The transmembrane currents of the channels *I*_K_, *I*_Na_ and the leakage current *I_L_*, which are assumed outward, read

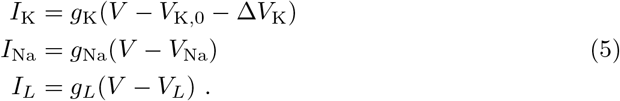

Here, 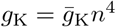 and 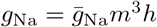. The terms 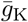 and 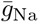 are the channel densities, *g_L_* is the membrane conductance, *V*_Na_ is the Na^+^ reversal potential. *V_L_* sets the rest potential *V_r_* of the membrane at Δ*V*_K_ = 0 and *I_syn_* = 0.

In our model, we assume the presence of ionic actuators as the sole responsible for the perturbation of the extracellular concentration of K^+^, thereby adopting *V*_K_ as a control input of the model. However, there exist several physiological actors that are involved in the regulation of K^+^ concentration and in principle may interfere with the action of the ionic actuators. Examples include the Na^+^/K^+^ pumps, the buffering of glial cells, or diffusion in the extracellular microenvironment [15,36]. Our choice of neglecting such phenomena is prompted by recent advances in the technology of ionic actuators, which is reaching temporal resolutions comparable to the time scales of synaptic activity [37, 38]. From a mathematical perspective, we perform a time scale separation [12,39] that disentangles the fast dynamics of action potentials generation (~ milliseconds) and ionic actuators operation (~ tens of milliseconds) from the slower dynamics of the regulation of ionic concentrations (~ seconds).

We support our discussion with analyses of several HH-type models taken from the literature, which are calibrated on electrophysiological data of cortical neurons in rats. As a comparison, we also consider the classical HH model of the squid giant axon. All the key references of these models are summarized in Table 1, while the details regarding their membrane parameters and channel kinetics can be found in Appendix A. Some of these models slightly differ from the model in Eq. (4) that is considered for the mathematical derivations in the sections Equilibrium points of the model and The conductance of K^+^-selective channels affects switchings of the class of neural excitability in the Δ*V*_K_ – *I_syn_* plane. For instance, they may include further voltage-gated channels or K^+^ and Na^+^ specific leak currents. The generalization of such derivations to these cases can be found in Appendix B.

**Table 1.** HH-type models used as case studies. With the exception of the classical model of the squid giant axon, all the models have been fitted as single-compartment models to reproduce electrophysiological measurements of rat cortical neurons. We refer to Appendix A for the details regarding the membrane parameters and the channel kinetics.

| Codename | Description of the model | Reference |
| --- | --- | --- |
| squid-hh52 | Original HH model for the squid giant axon. | [34] |
| rat-wei14 | Excitatory pyramidal neuron from the CA1 region of the hippocampus. | [14, Methods] |
| rat-cressman09 | Inhibitory interneuron from the CA1 region of the hippocampus. | [11, Methods] |
| rat-wang96 | Fast-spiking inhibitory interneuron from the CA1 region of the hippocampus. | [40, Methods] |
| rat-pospischil08-FSinh | Fast-spiking inhibitory interneuron from the somatosensory cortex. | [41, Fig. 4] |
| rat-pospischil08-RSexc | Regular-spiking excitatory pyramidal neuron from the somatosensory cortex. | [41, Fig. 2a] |

### Equilibrium points of the model

We deploy bifurcation analysis [32, 33] to investigate the qualitative behavior of the system (4) when both ionic and current stimuli are present. In this regard, the equilibrium points (also denoted as steady states) play a central role. They are found imposing the time derivatives in Eqs. (4) to zero, and therefore constitute a vector in the form (*n*_∞_(*V_s_*), *m*_∞_(*V_s_*), *h*_∞_(*V_s_*), *V_s_*). The allowed values of the membrane potential at equilibrium *V_s_* are set by Δ*V*_K_ and *I_syn_* according to

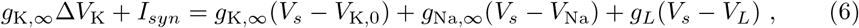

where *g*_K,∞_, *g*_Na,∞_ are the steady-state conductances, evaluated at *x* = *x*_∞_(*V_s_*), with *x* ∈ {*n*, *m, h*}. Since the steady-state variables are solely a function of *V_s_*, we shall denote an equilibrium point by only the value of *V_s_*. We point out that, for a fixed *I_syn_* (respectively, Δ*V*_K_), Eq. 6 establishes a map that provides the value of Δ*V*_K_ (respectively, *I_syn_*) required to set a given *V_s_*. Namely,

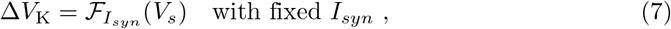

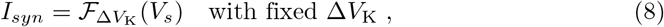

where we have defined the functions

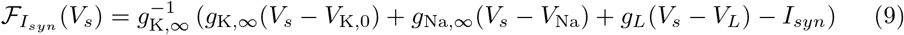

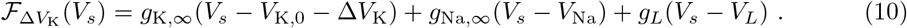

The stability of an equilibrium point *V_s_* can be assessed by studying the Jacobian matrix *J* of the system, with respect to the state variables (*n, m, h, V*), evaluated in such point:

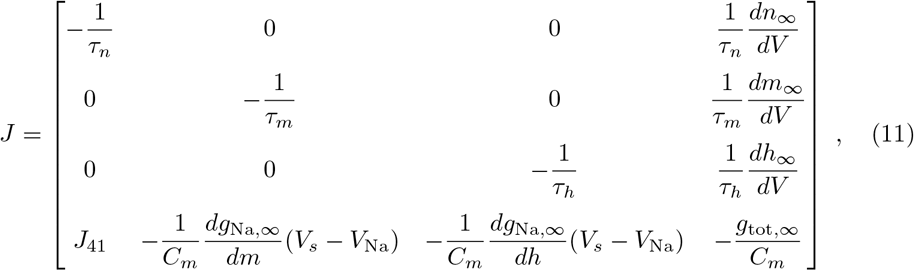

with *g*_∞,*tot*_ = *g*_K,∞_ + *g*_Na,∞_ + *g_L_* and *J*_41_ the only entry that explicitly depends on Δ*V*_K_,

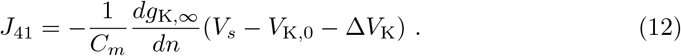

Equivalently, *J*_41_ can be expressed as a function of *I_syn_* as

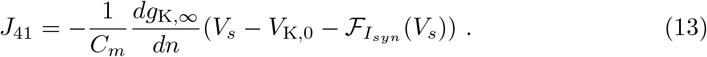

An equilibrium point *V_s_* is (asymptotically) stable if all the eigenvalues λ_*i*_ of *J* have a negative real part. Conversely, it is (exponentially) unstable if at least one eigenvalue λ_*i*_ of *J* has a positive real part [32]. Assuming Δ*V*_K_ as bifurcation parameter and *I_syn_* fixed, we can study the stability of the equilibrium points *V_s_* inspecting the eigenvalues traces λ_*i*_(*V_s_*) of the Jacobian matrix (11), using *J*_41_ as in Eq. (13). Equivalently, *I_syn_* can be used as bifurcation parameter, fixing Δ*V*_K_ and using *J*_41_ as in Eq. (13) in the Jacobian. We used such methodology in the study of bifurcations of equilibria discussed in the Results section, which was carried out in Matlab [42]. We also studied the bifurcations of limit cycles resorting to the MatCont toolbox [43].

### Two-compartment model

The single-compartment model of Fig. 1 mimics a setting where the ionic perturbation occurs throughout the microenvironment surrounding the neuron. However, the spatial resolution of ionic actuators has already approached the single-cellular and sub-cellular scale of micrometers, paving the way to an ionic actuation that is spatially localized at the neural membrane, e.g., to a portion of the soma. As a result, the perturbation of K^+^ extracellular concentration directly affects only the K^+^ channels in such membrane patch. To account for such a scenario, we also consider the two-compartment model in Fig. 2. It is inspired by the Pinsky-Rinzel model [44], originally introduced to lump the interactions between soma and dendrites. The dynamics of the gating variables in each compartment are ruled as in Eq. (4). Instead, the current balance equations in the two compartments now include a coupling term. Namely,

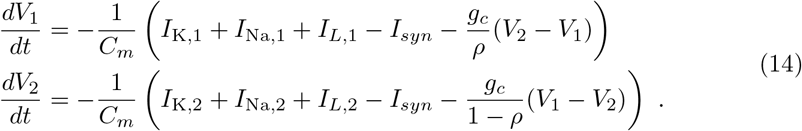

The first compartment lumps the dynamics of the actuated patch of the membrane, i.e., that directly perceives the perturbation of the [K^+^]. The second compartment refers instead to the non-actuated patch. *ρ* represents the fraction of membrane area associated with the actuated patch, while *g_c_* is a conductance per unit area that accounts for the electrical coupling between the actuated and non-actuated patches. They both mimic the influence of the geometry in the problem. The currents of the channels in the two compartments are

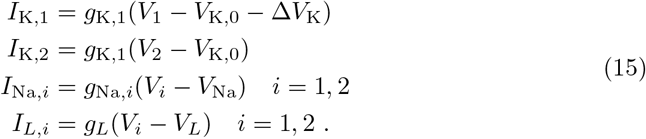

We point out that Δ*V*_K_ is applied only to the first compartment, while *I_syn_* is applied equally to both compartments. We use this Pinsky-Rinzel-like model to investigate how the fraction of ionically actuated membrane affects the proposed neuromodulation strategy in the last part of the Results section. To support our discussion, we still employ the HH-type models listed in Table 1, whose membrane parameters and channel kinetics are now associated with both the two compartments.

**Fig 2.**
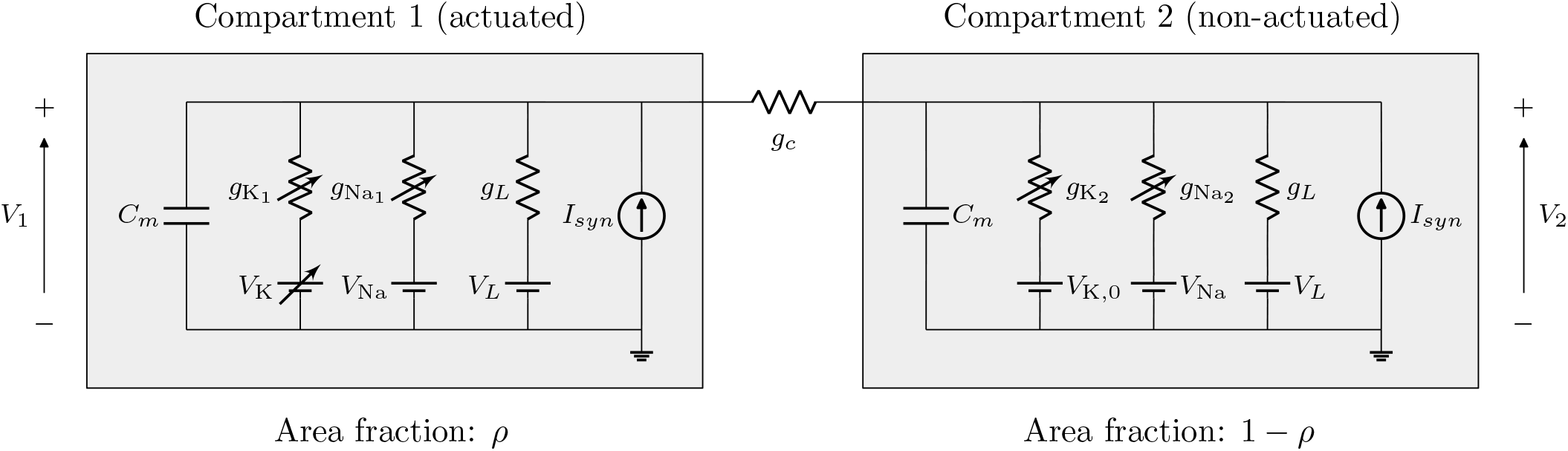
Two-compartment model. It mimics a scenario where an ionic actuator of sub-cellular size is employed. The actuated (respectively, non-actuated) compartment lumps the dynamics of the membrane patch that perceives (respectively, does not perceive) the action of the ionic actuator through the manipulation of [K^+^], and hence the variation of *V*_K_. The same current *I_syn_* is assumed in both compartments. The parameter *g_c_* is a conductance per unit of area and accounts for the electrical coupling between the two regions. The currents in the first (respectively, second) compartment are associated to the membrane area *A*_1_ (respectively, *A*_2_), while the current across *g_c_* is associated to the total membrane area *A_tot_* = *A*_1_ + *A*_2_. As a consequence, the current balance equations of the two compartments are as in (14), with *ρ* = *A*_1_/*A_tot_* and 1 – *ρ* = *A*_2_/*A_tot_*.

## Results

In the present section, we first leverage methods of bifurcation analysis to understand the impact of *I_syn_* and Δ*V*_K_ on neural excitability. We then exploit these findings to build a principled approach to neuromodulation via ionic actuation of potassium. Further, we provide a motivation for the switchings of neural excitability in the *I_syn_ –* Δ*V*_K_ plane, and we investigate the effects of a spatially localized actuation.

### Bifurcation analysis of the system

A typical neuron may exhibit three steady regimes: rest, spiking, and depolarization block (in the following, simply block). At rest and block, it eventually attains a stable value of membrane potential. It is more depolarized in the latter case than in the former. While spiking, the neuron generates a repetitive train of action potentials. If a steady regime is the only one possible, we refer to it as *tonic*. Otherwise, if multiple such regimes coexist, it is possible for the system to switch between them, and therefore, we refer to it as *bistable*. We point out that with spiking, we herein refer to the presence of a generic periodic trajectory, disregarding possible classifications according to the firing patterns, such as the distinction between regular spiking, fast spiking, or intrinsically bursting neurons [45].

In a neuromodulation scenario, it is pivotal to understand which regimes arise in the space of the input variables. Indeed, this enables a principled framework to effectively steer the system to the desired behavior through the controllable input to the model (e.g., Δ*V*_K_), possibly in a robust fashion with respect to uncontrollable input (e.g., *I_syn_*). In addition, the identification of regions of tonic behavior avoids undesired switching of the system between bistable configurations, which could result from the presence of noise or fluctuations of the model parameters. To these aims, we can deploy methods of bifurcation analysis since the aforementioned neural behaviors readily translate into properties of equilibrium points and stable limit cycles [32]. Hereafter, we first study the qualitative behavior and bifurcations of the system (4) with respect to a single input parameter (either Δ*V*_K_ or *I_syn_*), while the other is kept fixed. We then show how the analysis generalizes to the Δ*V*_K_ – *I_syn_* plane. For ease of discussion, we describe the qualitative changes of the system’s dynamics via bifurcations as observed while increasing either Δ*V*_K_ or *I_syn_*. Therefore, with a little abuse of language, we refer to a qualitative behavior, or a bifurcation, as occurring *before*, or *after* one another, as we implicitly assume to move along the axis of one of these two parameters.

The bifurcations of equilibrium points are of two kinds: saddle-node and Hopf. In a saddle-node bifurcation, a stable and an unstable equilibrium coalesce and cancel each other. In a Hopf bifurcation, a stable equilibrium loses or gains stability. To identify such bifurcations, we consider the eigenvalue traces λ_*i*_ of the Jacobian matrix of the system (11), as introduced in the Methods section. We define *V_th_* (respectively, *V_block_*) as the value of *V_s_* where the eigenvalue traces enter into instability (respectively, recover from instability), see Fig. 3 (left). At *V_th_* and *V_block_*, bifurcations of equilibrium points occur. They can be revealed through the bifurcation diagrams as in Fig. 3 (right), namely plotting *V_s_* as a function of the bifurcation parameter through Eq. (7) or (8). *V_th_* corresponds to a saddle-node bifurcation if therein a real eigenvalue changes sign from negative to positive. Instead, *V_th_* corresponds to a Hopf bifurcation if therein the real parts of a pair of complex conjugate eigenvalues change sign from negative to positive. *V_block_* typically corresponds to a Hopf bifurcation since it occurs in a monotonically increasing interval of the *V_s_* curve [32]. Therefore, therein the real parts of a pair of complex conjugate eigenvalues change sign from positive to negative. For the sake of simplicity, we shall use *V_th_* and *V_block_* to refer to both the equilibrium points where such bifurcations occur, as well as to the bifurcations themselves. The rest regime disappears at *V_th_*, which therefore acts as a threshold for the system towards a spiking trajectory (when present, see below). Conversely, the block regime appears after *V_block_*, thereby enabling the system to escape from a spiking trajectory. Fig. 3i shows that, if *V_th_* is a saddle-node, it may take place before or after *V_block_*, depending on the value of the input parameter kept fixed during the analysis (*I_syn_* in the figure). In the former case, the system shows only an unstable equilibrium between the two bifurcations, and therefore a tonic spiking regime is allowed in such interval. In the latter case, the system is bistable with two stable equilibria between the two bifurcations, i.e., the rest and block configurations, and the possibility of eliciting a tonic spiking behavior is ruled out. Even though the notion of *bistability* is usually employed in the literature to refer to the generic coexistence of stable trajectories [32], either equilibrium points or limit cycles, we herein restrict this notion to such coexistence of rest and block configurations.

**Fig 3.**
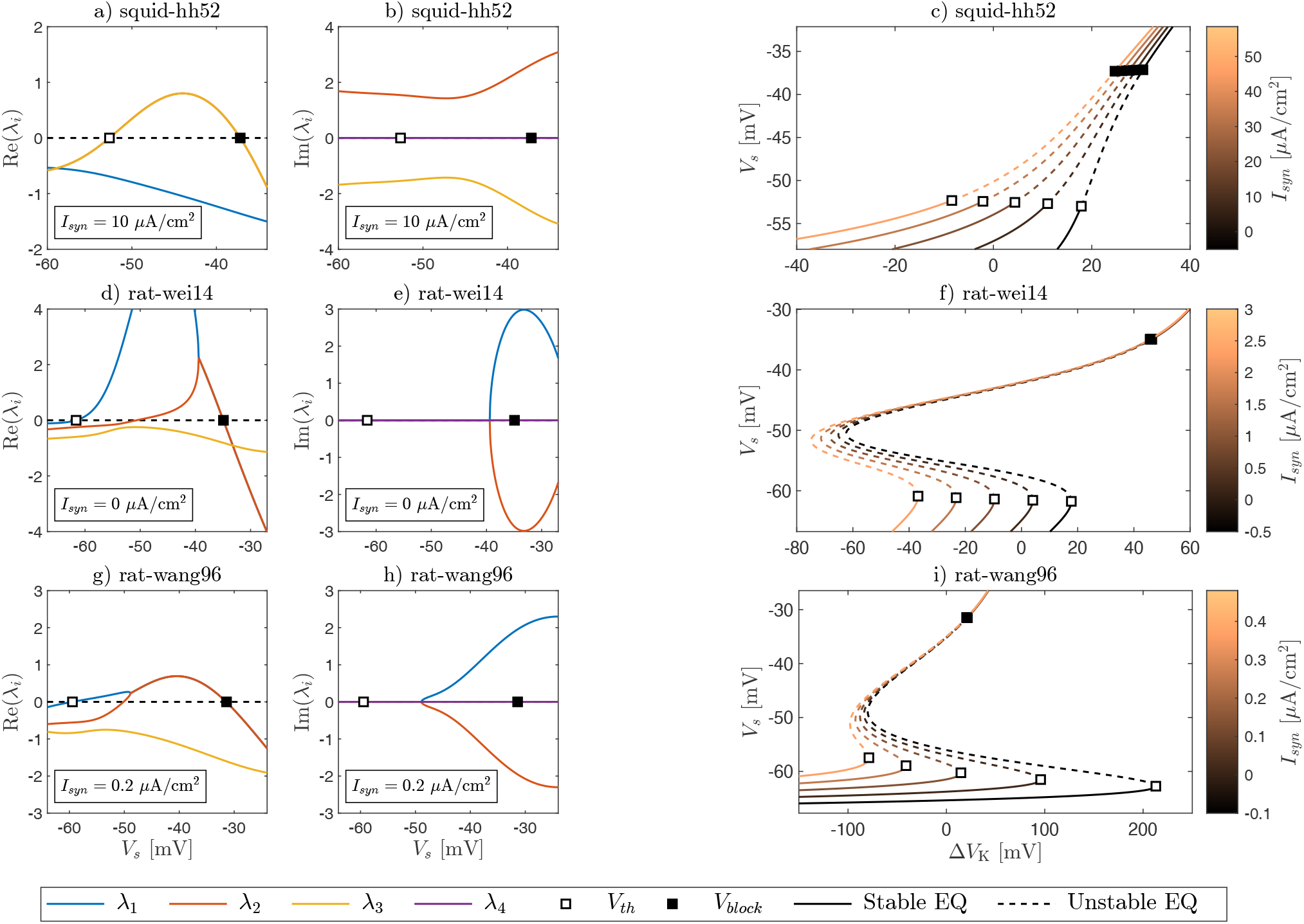
Bifurcation analysis of equilibrium points. The equilibrium points (EQ) of the system (4), along with their stability properties, are studied as described in the section Methods. Δ*V*_K_ is assumed as bifurcation parameter. The same plots are reported for the HH-type models squid-hh52, rat-wei14, rat-wang96 (see Table 1). Left) Real and imaginary parts of the eigenvalue traces λ_*i*_ of the Jacobian matrix (11) as a function of the equilibrium value of the membrane potential *V_s_*, with fixed *I_syn_* (see insets). These curves enable us to distinguish between stable and unstable *V_s_* by inspecting the sign of the real parts of the eigenvalues. Right) Bifurcation diagrams of the equilibrium points obtained plotting the equilibria *V_s_* as a function of Δ*V*_K_ according to Eq. (7). These diagrams are shown for different choices of *I_syn_* and augmented with the stability information extracted from the study of the eigenvalue traces. *V*_th_ may correspond to a Hopf (a,b,c), or a saddle-node (d-i), bifurcation, while *V_block_* is associated to a Hopf bifurcation. (i) In the case of a saddle-node at *V_th_*, the bifurcation can occur before or after *V_block_* in terms of Δ*V*_K_, depending of the value of *I_syn_*.

The study of bifurcations of stable limit cycles is in general more involved. However, for our concerns, the bifurcations that give rise to stable limit cycles are of three kinds: fold limit cycle, saddle-node on invariant circle (SNIC), and saddle-homoclinic (SH) [32]. Fig. 4 shows some examples of bifurcation diagrams of both limit cycles and equilibria for the models under study. A fold occurs before *V_th_* when it is Hopf subcritical, and results in a pair of stable and unstable limit cycles. A SNIC occurs at *V_th_* when it is a saddle-node but the system is not bistable. An SH bifurcation occurs before *V_block_* when the system is bistable. Eventually, such stable limit cycles disappear, either through a limit cycle bifurcation slightly after *V_block_*, if it is Hopf subcritical, or at *V_block_*, if it is Hopf supercritical. Unstable limit cycles also either appear or disappear at fold limit cycle bifurcations. However, such trajectories are not of practical concern due to their repulsive nature. Figs. 4h,i confirm that the bifurcation *V_th_* must occur before *V_block_* to elicit tonic spiking. Non-tonic spiking is allowed when fold or SH bifurcations are involved. In such cases, limit cycles may coexist with rest (Figs. 4a,b,c,h), block (Fig. 4i) or bistable (Fig. 4h) regimes. In these cases, a tonic rest (respectively, block) regime is reached sufficiently far from *V_th_* (respectively, *V_block_*).

**Fig 4.**
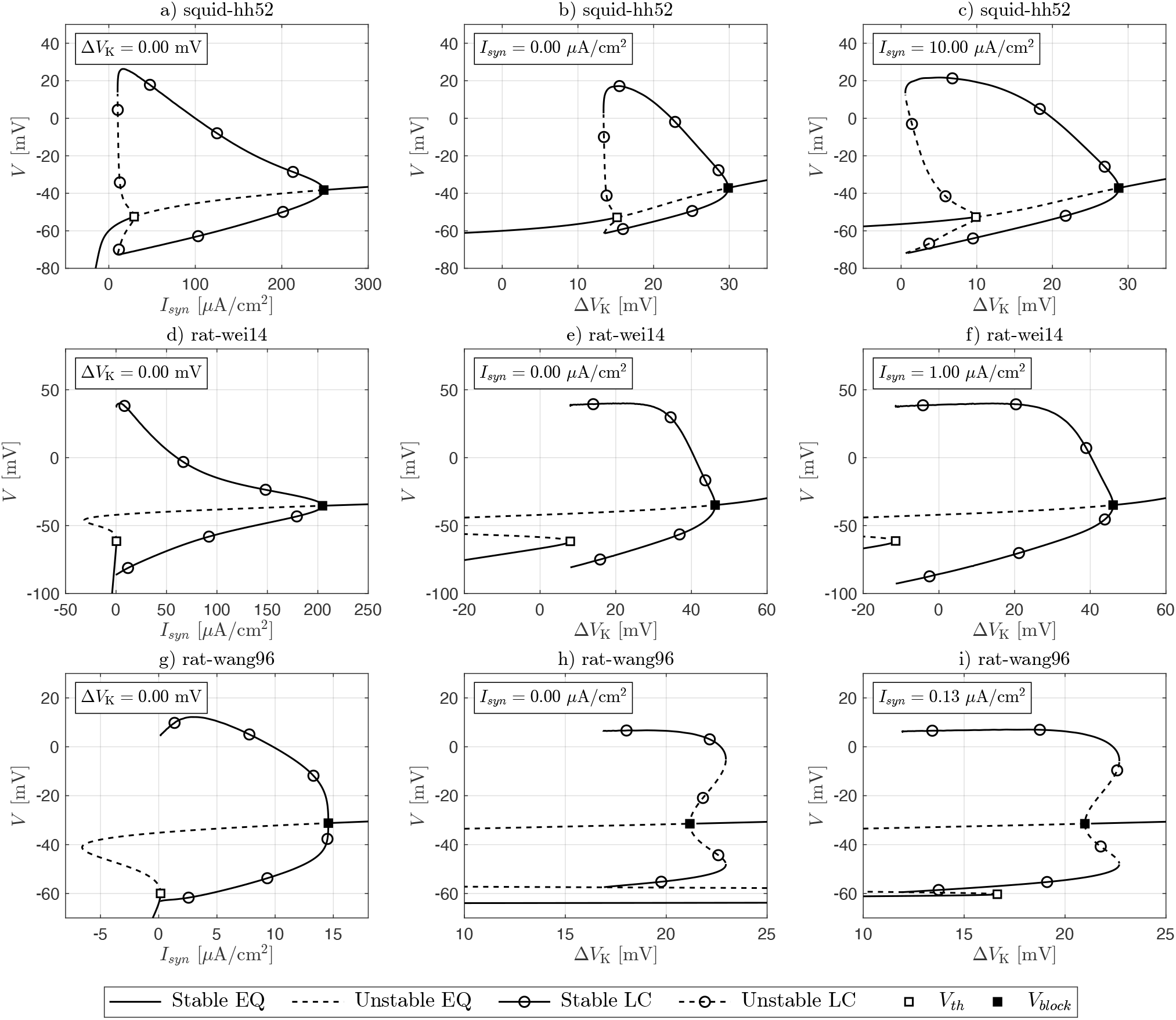
Bifurcation analysis of limit cycles. Some representative bifurcation diagrams for the different HH-type models adopted as case studies (see Table 1) are shown. They include both the limit cycles (LC) and the equilibrium points (EQ), along with their stability properties. Limit cycles are obtained using the MatCont [43], and are represented through their maximum and minimum values of membrane potential. Equilibria are studied as in Fig. 3. a,d,g) The first column refers to the case of purely current actuation, where *I_syn_* is used as the bifurcation parameter and Δ*V*_K_ = 0. The bifurcation *V_th_* occurs before *V_block_* and a tonic spiking regime can exist in between. The stable limit cycle arises either with (a) a fold limit cycle before *V_th_* or (d,g) a SNIC at *V_th_*. It disappears with a supercritical Hopf bifurcation at *V_block_*. Before *V_th_*, a rest regime is possible, which is always tonic in the SNIC case and only before the fold. After *V_block_*, the system enters into a tonic block regime. b,e,h) The second column refers to the case of purely ionic actuation, with Δ*V*_K_ as the bifurcation parameter and *I_syn_* = 0. In (b,e) the models retain the same qualitative behavior as in the case of current actuation. Conversely, in (h), *V_block_* occurs before *V_th_* and therefore a tonic spiking behavior is not possible (*V_th_* = –61.7 mV and Δ*V*_K,*th*_ = 110 mV are not shown because out of scale). Instead, the system is bistable between these two bifurcations, with both rest and block behaviors accessible. Interestingly, also a stable limit cycle exists, which is not tonic. It arises via an SH bifurcation and eventually disappears in a fold. c,f,i) The third column considers the case of ionic actuation in the presence of an excitatory current *I_syn_*. As a result, a lower Δ*V*_K_ is sufficient to give rise to a stable limit cycle in (c,f) when compared to the cases in (b,e). Further, the bistable region may disappear, thereby making tonic spiking possible. Compare for instance (i) and (h). Interestingly, the bifurcations that undergo equilibria and limit cycles may change depending on whether *I_syn_* or Δ*V*_K_ is used as the bifurcation parameter. For instance, compare the disappearance of the stable limit cycle via Hopf supercritical in (g) and a fold in (h,i).

Neurons may be classified into three classes of neural excitability depending on the properties of their f-I curve, i.e., the steady-state relation between the input current and the firing frequency. Class I (respectively, II) neurons show a continuous (respectively, discontinuous) f-I curve, while in class III neurons no firing is elicited irrespective of the applied current [32]. Bifurcation diagrams allow us to infer the excitability class of a neuron, and we refer the reader to [32] for further details. From the previous analyses follows that, when no ionic actuation is present (Δ*V*_K_ = 0), squid-hh52 shows class II excitability (see Fig. 4a), while rat-wei14 and rat-wang96 show class I excitability (see Figs. 4d,g). We can introduce an analogous classification assuming Δ*V*_K_ as input of the model. In such setting, squid-hh52 and rat-wei14 maintain the same excitability type (see Figs. 4b,c,e,f). Conversely, rat-wang96 shows class I excitability when a sufficiently large *I_syn_* is applied (not shown), but it switches to class II when *I_syn_* is in a certain range around 0.13 *μ*A/cm^2^ (see Fig. 4i). This happens because an SH bifurcation gives rise to the limit cycle instead of a SNIC. For smaller values of *I_syn_*, rat-wang96 further switches to class III excitability (see Fig. 4h) due to the disappearance of tonic spiking and the appearance of bistability. The possibility of a Δ*V*_K_-induced switching from class I to class II has been recently pointed out in the literature [39] and corroborated *in-vitro*. Our analysis complements these findings, suggesting that these switchings may occur or not depending on the type of neuron considered. Further, our study emphasizes the possible emergence of a class III excitability when an ionic actuator drives the system without a sufficiently large synaptic input. This fact may be detrimental from a neural engineering perspective since it rules out the possibility of eliciting tonic spiking in certain types of neurons.

In the previous analyses, we considered only one input variable among Δ*V*_K_ and *I_syn_* as a bifurcation parameter while keeping the other fixed. Repeating the studies for different values of the fixed parameter, it is thus possible to find the boundary curves Δ*V*_K,*th*_ – *I_th_* and Δ*V*_K,*block*_ – *I_block_* that identify where the bifurcations *V_th_* and *V_block_* take place in the space of input variables. This leads us to the two-dimensional bifurcation diagrams as in Fig. 5. They contain four regions: *rest, spike, block*, and *bistable*. In the *spike* region, the system already underwent the *V_th_* bifurcation, but not yet *V_block_*. Models in this region exhibit tonic spiking behavior. In the *rest, block*, and *bistable* regions, the homonym regimes are reachable, but may also coexist with limit cycles, as previously observed. Such limit cycles typically disappear sufficiently close to the boundary curves. An exception in this regard is the squid-hh52 model, where the fold bifurcation occurs at large (negative) Δ*V*_K_ if a rather large excitatory current *I_syn_* is present, and hence far away from the Δ*V*_K,*th*_ – *I_th_* curve (not shown). Nevertheless, we hereafter focus on the other models employed as case studies (see Table 1, since cortical neurons are of major concern as targets in neuromodulation scenarios. Consequently, the bifurcation analysis of equilibria suffices to establish a principled framework to effectively drive the system to the desired tonic regime via ionic actuation, as we shall see in the next section. We point out that the possibility of neglecting the bifurcation analysis of limit cycles reduce considerably the computational burden of the study. Fig. 5 further confirms that, depending on the neuron model considered, the transition from rest to (tonic) spiking may not be possible acting on Δ*V*_K_ alone, without the presence of an excitatory current *I_syn_*. In Fig. 6, we corroborate the predictions of Fig. 5 for the model rat-wang96, showing the steady regimes reached by the model in each one of the four regions for different initial conditions.

**Fig 5.**
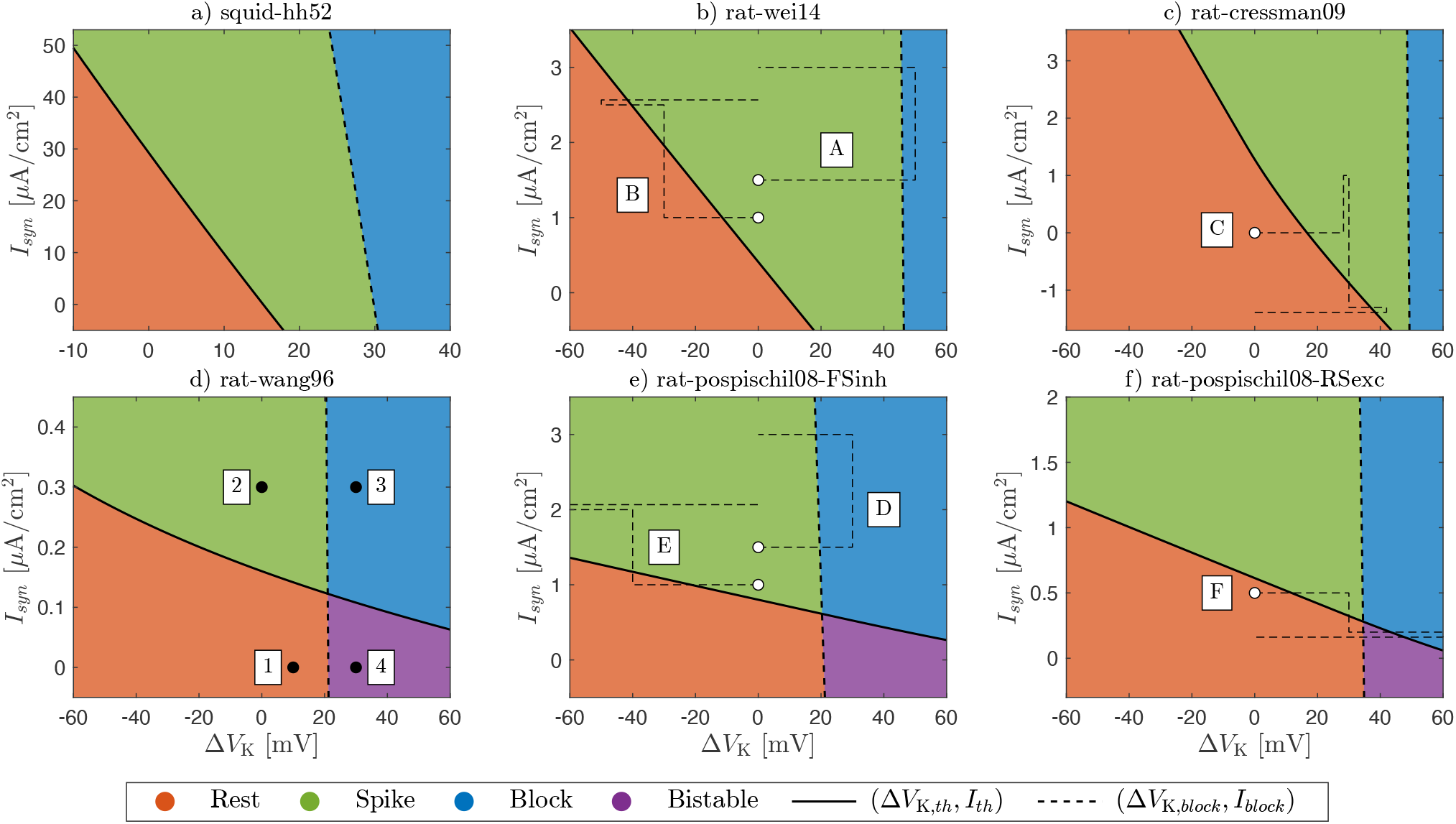
Two-dimensional bifurcation analysis of equilibrium points. The diagrams show the boundary curves Δ*V*_K,*th*_ – *I_th_* and Δ*V*_K,*block*_ – *I_block_*, which correspond to the occurrence of the *V_th_* and *V_block_* bifurcations in the space of input parameters Δ*V*_K_, *I_syn_*. The diagrams of all the HH-type models adopted as case studies (see Table 1) are shown. They are obtained by repeating the one-dimensional bifurcation analysis depicted in Fig. 3 for different values of the fixed parameter *I_syn_*. They distinguish the four regions *rest*, *spike*, *block*, and *bistable*, named after the regimes that can emerge therein. In *spike*, a tonic spiking is guaranteed, while, in the other regions, a spiking regime may coexist if limit cycle bifurcations as SH or fold take place. Therefore, to attain a tonic regime in *rest* and *block* is necessary to move sufficiently far away from the boundary curves. In *bistable* regions, both the rest and block regimes are allowed. The enumerated markers in (d) refer to the simulations in Fig. 6. The dashed paths labeled with uppercase letters in (b,c,e,f) refer to the example simulations of inhibition and stimulation of electrical activity in Figs. 7 and 8, with the marker denoting the initial condition.

**Fig 6.**
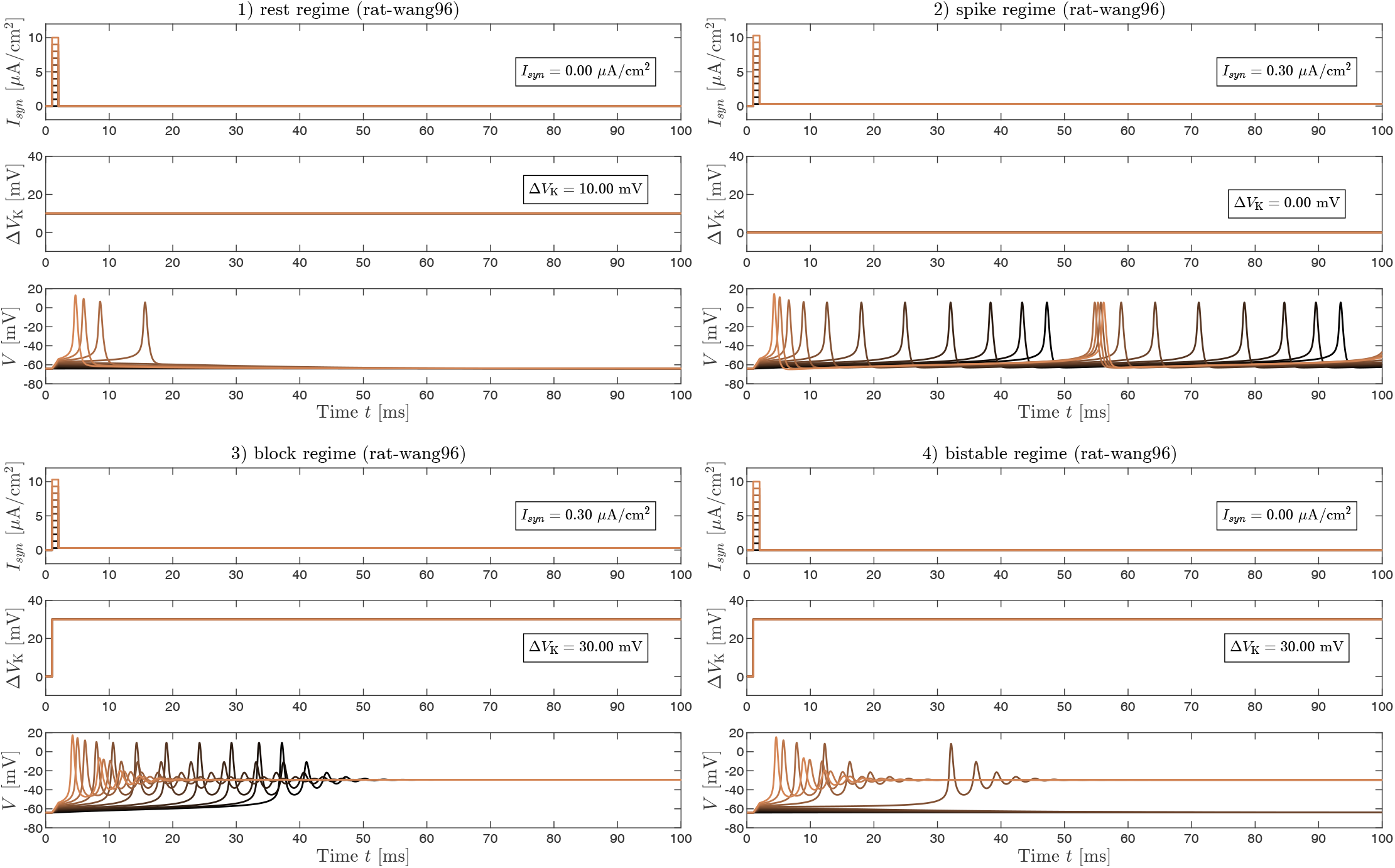
Possible tonic regimes and bistability. Simulations of the model rat-wang96 for four combinations of Δ*V*_K_ and *I_syn_* values, each one corresponding to a different region of Fig. 5d (see Fig. 5’s caption). Each simulation is repeated for several initial conditions (different colors), which are set by rapid current pulses at the beginning of the transients (duration: 1 ms; amplitude: between 1 and 10 *μ*A/cm^2^). In 1), 2), and 3), a rest, a spiking, and a block regime are achieved respectively, irrespective of the initial condition. In 4), the steady state may be either a rest or a block condition due to the bistable configuration of the system. These simulations corroborate the theoretical findings reported in Fig. 5. We point out that 1), 3), 4) are chosen sufficiently far from the boundaries to avoid the presence of a limit cycle. Compare e.g., Figs. 5d and 4h.

### Principled neuromodulation via ionic actuation

In the scenario of neuromodulation via ionic actuation under study, one aims to elicit the desired regime of electrical activity in a neuron via a controlled perturbation of the extracellular concentration of K^+^. In a stimulation application, the target is to trigger electrical activity in the neuron, hence a spiking regime. In an inhibition application, the target is to silence electrical activity in the neuron. This can be accomplished by either reaching a rest or a block configuration. The target regime is expected to be maintained robustly, namely regardless of possible synaptic currents, at least in a reasonable range of values. Further, the induced behavior should ideally be tonic to avoid undesired switches to other regimes due to noise or other disturbances.

The previous desiderata can be accomplished in a principled fashion by means of the theoretical framework developed in the previous section, especially resorting to the two-dimensional bifurcation diagrams in Fig. 5. Indeed, given a range of values for *I_syn_*, it is possible to assess the extent of ionic release Δ*V*_K_ > 0, or ionic uptake Δ*V*_K_ < 0, that would be required to reach the target regime. Regarding inhibition, we observe that the curves associated with the *V_block_* bifurcation have a dependence on *I_syn_* much weaker than the ones associated with *V_th_*. Therefore, the maintenance of a block regime is expected to be much less sensitive to the presence of a synaptic current than the maintenance of a rest regime. We point out that a block regime can be elicited also with pure current actuation (see Fig. 4a, d, g). It is not evident from Fig. 5 because it requires a rather unphysiological current amplitude.

Fig. 7 shows examples of inhibition of electrical activity via ionic actuation. As previously observed, a neuron driven in the block regime persists in such regime quite irrespective of the synaptic current applied. Furthermore, the minimum extent of Δ*V*_K_ needed to reach this regime depends on the model and hence the type of the neural cell considered, see Fig. 7a, b. If rest is assumed as a target, the maintenance of such behavior is much more sensitive to the value of *I_syn_* and may require a more accurate online adjustment of Δ*V*_K_. However, when an excitatory *I_syn_* is present, we observe that it rapidly may amount to a large (negative) Δ*V*_K_, see Fig. 7d. However, this implies the need for an almost complete depletion of extracellular K^+^ according to Eq. (3), which is expected to be hardly feasible in a realistic application.

**Fig 7.**
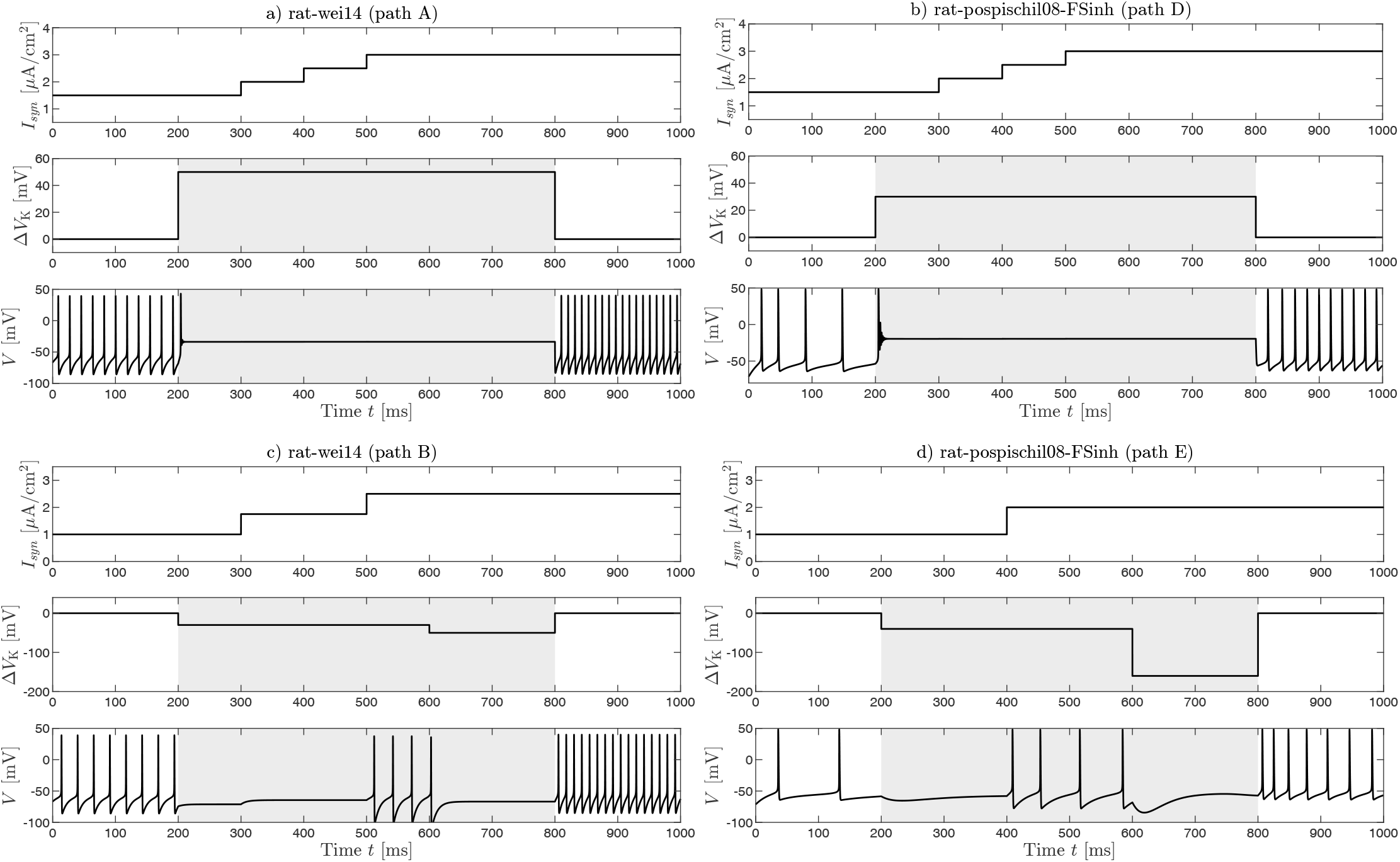
Examples of inhibition of electrical activity via ionic actuation. The shaded area in each plot highlights when ionic actuation is active. Each simulation refers to a dashed path in Fig. 5 denoted with an uppercase letter (A, B, D, or E). The values of Δ*V*_K_ were chosen as a function of *I_syn_* and the target regime. a,b) assume a block configuration as the target. The maintenance of such configuration is more robust to disturbance by external synaptic currents and requires a different minimum extent of Δ*V*_K_ to be established, depending on the model considered. c,d) assume a rest configuration as the target. In this case, the maintenance of the regime is much more sensitive to *I_syn_*, which may imply the need for a large Δ*V*_K_ < 0, as in d). Such situation could become rapidly unfeasible, leading to the complete depletion of [K^+^]_*o*_ (see Eq. (3)).

Fig. 8 shows examples of stimulation of electrical activity via ionic actuation. As previously observed, different models, and hence potentially different neuron types, may show or not a region of tonic spiking under purely ionic actuation. In the former case, tonic spiking be reached increasing Δ*V*_K_, see Fig. 8a. The minimum extent of Δ*V*_K_ needed to establish such a regime increases if a hyperpolarizing *I_syn_* is present. In the latter case, tonic spiking becomes reachable only if a sufficient excitatory *I_syn_* is present, see Fig. 8b. Otherwise, the tonic regimes the system can attain are only rest and block.

**Fig 8.**
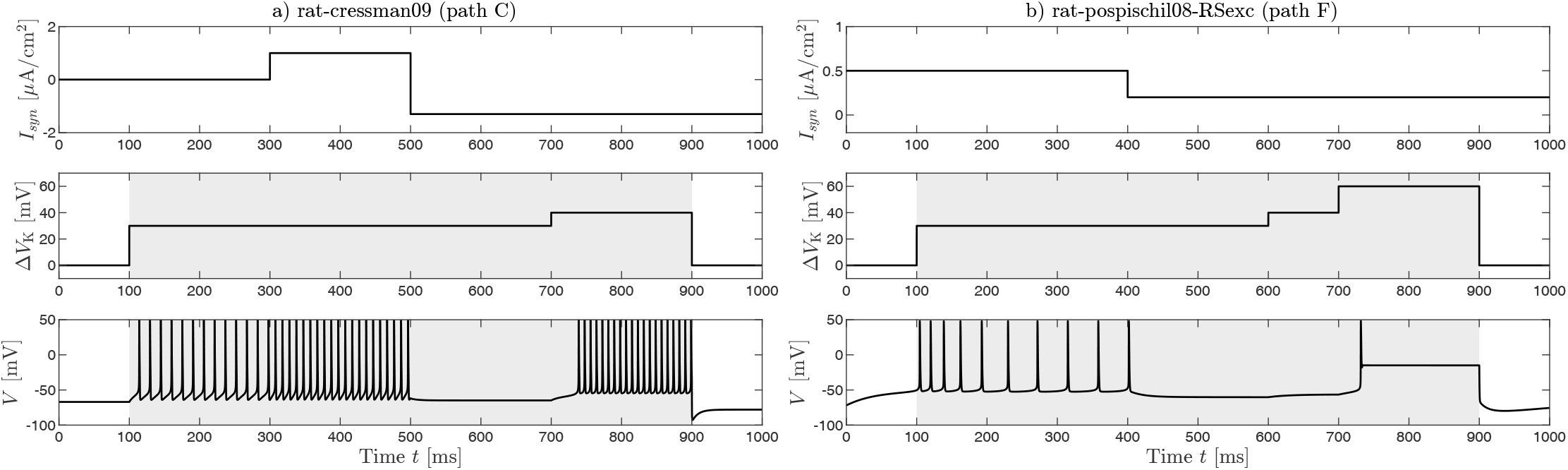
Examples of stimulation of electrical activity via ionic actuation. The shaded area in each plot highlights when ionic actuation is active. Each simulation refers to a dashed path in Fig. 5, denoted with an uppercase letter (C or F). The values of Δ*V*_K_ are chosen as a function of *I_syn_* and of the target regime. a) In this model, a regime of tonic spiking can be elicited through solely Δ*V*_K_. However, the minimum Δ*V*_K_ required increases if a hyperpolarizing *I_syn_* is present. b) This model does not show a region of tonic spiking in the case of purely ionic actuation. Such regime is therefore accomplished only in presence of a sufficiently large excitatory *I_syn_*. Otherwise, the possible tonic regimes are only rest and block.

### The conductance of K^+^-selective channels affects switchings of the class of neural excitability in the Δ*V*_K_ – *I_syn_* plane

From the previous analyses, it emerges that the qualitative dynamics of a neuron may differ depending on whether a current or an ionic actuator is used to drive the system. For instance, in the latter case tonic spiking may not occur. In other words, switchings of the class of neural excitability may occur in the *I_syn_ –* Δ*V*_K_ plane. An insight into such qualitative changes can be obtained by inspecting how the input variables locally affect the equilibrium points of the system (4). Namely, assuming a perturbation of the input variables Δ*V*_K_ → Δ*V*_K_ + *δV*_K_ and *I_syn_* → *I_syn_* + *δI_syn_*, a new equilibrium is established *V_s_* → *V_s_* + *δV_s_*. We can approximate *δV_s_* at the first-order as

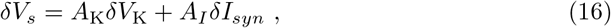

where *A_I_* is obtained by linearizing Eq. (6) with respect to *V_s_* and taking its inverse:

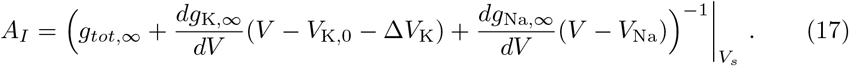

As a remark, *A_I_* is a function of both Δ*V*_K_ and *I_syn_* at the linearization point, with the latter dependence given implicitly by *V_s_*. The analogous definition for *A*_K_ yields

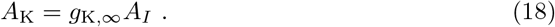

We observe that the transfer function of the ionic input *A*_K_ is further filtered by the conductance of the K^+^ channel. From this fact, natural comparisons with the current actuation scenario follow. First, in a rest state, i.e., *V_s_* < *V_th_*, the *g*_K_ is typically very small, and therefore a larger Δ*V*_K_ may be required to reach *V_th_* in order to leave the rest regime. Conversely, in a depolarization condition, i.e., *V* > *V_th_*, the *g*_K_ is much larger due to the non-inactivation of the K^+^ channels that shape the action potentials. Therefore, a smaller Δ*V*_K_ suffices to reach *V_block_* in order to establish a block regime. The impact of *g*_K_ may be so critical that Δ*V*_K,*th*_ overcomes Δ*V*_K,*block*_ and bistability replaces tonic spiking (see Fig. 4h). These observations are corroborated in Table 2, where *A_I_*, *A*_K_, *g*_K,∞_ are reported for the models under study, assuming the case of no ionic and current actuation as linearization point. We observe that *g*_K,∞_ is more than one order of magnitude smaller in the models that do not show tonic spiking in the case of purely ionic actuation (rat-wang96, rat-pospischil08-FSinh, rat-pospischil08-RSexc) with respect to the ones that show tonic spiking (squid-hh52, rat-wei14, rat-cressman09). Table 2 further reports the values of *I_syn_* and Δ[K^+^]_*o*_, where the bifurcations *V_th_* and *V_block_* occur in the case of either sole current or sole ionic actuation, respectively. This shows that, when tonic spiking is possible, the interval where it can be established in the space of the input parameters is wider in the latter case than in the former.

**Table 2.**
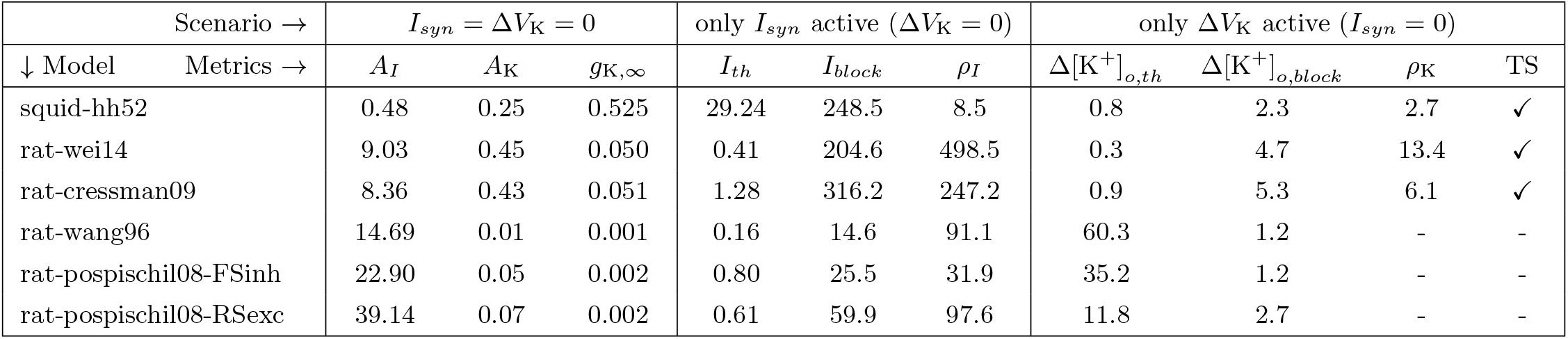
Comparison between ionic and current actuation. The *I_syn_* = Δ*V*_K_ = 0 scenario corresponds to an isolated neuron, with no supplied input. *A_I_* and *A*_K_ measure the attitude of *I_syn_* and Δ*V*_K_ to induce a perturbation of the steady state *V_s_*, see Eq. (16). They are related by the conductance of the K^+^ channel according to Eq. (18). The Δ*V*_K_ = 0 and *I_syn_* = 0 scenarios correspond to the cases of sole current and sole ionic actuation, respectively. Here, the values of the input where the bifurcations *V_th_* and *V_block_* occur are reported, with Δ[K^+^] in place of Δ*V*_K_ for a more physical comparison (computed with Eq. (3)). The ratios *ρ_I_* = *I_block_/I_th_* and *ρ*_K_ = Δ[K^+^]_*o*, *block*_/Δ[K^+^]_*o*, *th*_ measure the extension of the region of tonic spiking in the space of the input variable. The last column identifies if the model exhibits tonic spiking (TS) under sole ionic actuation. The metrics are evaluated considering the single-compartment model of Fig. 1. Units: *A_I_* is in (mS/cm^2^)^−1^, *g*_K,∞_ is in mS/cm^2^, *I_th_* and *I_block_* are in *μ*A/cm^2^, Δ[K^+^]_*o,th*_ and Δ[K^+^]_*o,block*_ are normalized to [K^+^]_*o*_.

### Influence of the fraction of actuated membrane

When an ionic actuator with a sub-cellular size is employed to modulate the electrical activity of a neuron, a two-compartment model as in Fig. 2 can be used to account for the local perturbation of *V*_K_ along the neural membrane. In this scenario, the impact of the ionic actuator on the membrane potential is expected to scale with the fraction of actuated membrane *ρ*, being that the K^+^ channels are directly recruited by Δ*V*_K_ only therein. Indeed, in the case of full coupling between the two compartments (*g_c_* → ∞), the current balance Eqs. (14) reduce to the single-compartment case of Eq. (4), with the only difference that the K^+^ current now reads

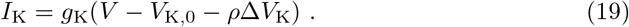

The single-compartment model considered in the previous sections is therefore retrieved assuming full coupling and *ρ* = 1. Consequently, if a value of Δ*V*_K_ enables to reach a certain regime according to Fig. 5, it must be increased by a gain factor *ρ*^−1^ to elicit the same result in the two-compartment model with full coupling but *ρ* < 1. The presence of a finite coupling *g_c_* between the two compartments is expected to alter such gain factor, possibly introducing new steady regimes in the system.

Fig. 9 shows examples of ionic inhibition and stimulation of electrical activity in the two-compartment case. The combinations of Δ*V*_K_ and *I_syn_* that elicit the target regimes can be compared with the ones predicted in Fig. 5 for the single-compartment case. We observe that to accomplish inhibition the minimum required Δ*V*_K_ is increased by more than a *ρ*^−1^ factor (compare Figs. 9a, b with Figs. 5e, b). Conversely, to obtain a similar behavior in the stimulation setting is sufficient a factor lower than *ρ*^−1^ (compare Figs. 9c and 5c). Interestingly, certain models reveal new steady regimes in which the actuated membrane patch is in block while the non-actuated one is spiking (see Fig. 9d). However, such behavior can be silenced by increasing Δ*V*_K_ (not shown).

**Fig 9.**
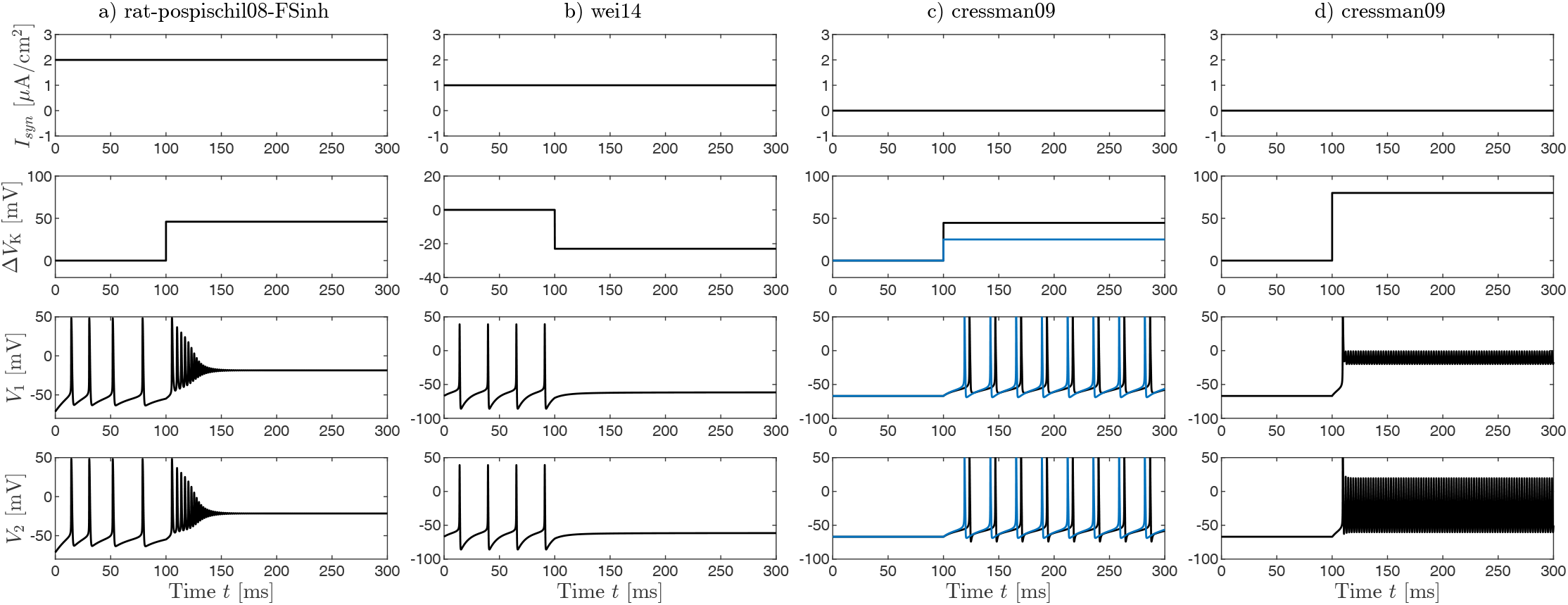
Examples of neuromodulation with ionic actuators of sub-cellular size. The two-compartment model in Fig. 2 is simulated assuming half of the cell membrane is actuated (*ρ* = 0.5) and a coupling of *g_c_* = 2 mS/cm^2^ (black traces). a) block regime elicited with Δ*V*_K_ = 46 mV. b) rest regime elicited with Δ*V*_K_ = –23 mV. In a) and b), the reported Δ*V*_K_ is the minimum required to induce the target regime, rounded to the units digit. c) spiking regime elicited with Δ*V*_K_ = 44.6 mV, chosen to have the same spiking frequency obtained in the single-compartment model with Δ*V*_K_ = 25 mV (blue traces). d) mixed regime elicited with Δ*V*_K_ = 80 mV, where the actuated patch is in block while the non-actuated one is in (very fast) spiking. The oscillations in the first compartment are a consequence of the electrical coupling with the second compartment.

## Discussion

We considered reduced single-compartment HH models to investigate the possibilities and limits offered by a neuromodulation mechanism based on the perturbation of the extracellular K^+^ concentration. For the sake of a comparative study, we took into account both the classical model of the squid giant axon, along with five models of rat cortical neurons, see Table 1. We augmented them with the action of an ionic actuator, affecting the neuron via variations of the potassium reversal potential Δ*V*_K_, and with an external synaptic current *I_syn_*.

We first leveraged bifurcation analysis to characterize the phenomenology of the neural electrical activity in terms of the extent of ionic actuation Δ*V*_K_ and the action of synaptic current *I_syn_*. This enabled us to identify four regions in the Δ*V*_K_ – *I_syn_* plane. Namely, a region where tonic spiking can be elicited, two regions where rest and block regimes are allowed and can be tonic, and a bistable region where the system can be switched between rest and block states. Spiking behaviors might be possible not only in the homonym region of the space of bifurcation parameters, but they are not tonic and are confined nearby the boundary curves Δ*V*_K,*th*_ – *I_th_* and Δ*V*_K,*block*_ – *I_block_*. We showed that, in certain models, it is possible to elicit a tonic spiking behavior via ionic actuation only if an excitatory current of sufficient amplitude is present. Conversely, a bistable behavior manifests. We found that the main cause of this fact is a very small sub-threshold conductance of the K^+^ channels. Further, the extent of Δ*V*_K_ required to reach the block regime shows a weak dependence on *I_syn_*. The same does not apply to the rest regime since its maintenance in the presence of an excitatory current *I_syn_* may imply the need for an almost complete depletion of [K^+^]_*o*_.

These findings allow us to approach the neuromodulation task in a principled fashion. Indeed, for a given *I_syn_* and a target regime, it is possible to choose Δ*V*_K_ to effectively steer the system with the information provided by the two-dimensional bifurcation diagram in Fig. 5. We tested these predictions in simulations, showing examples of inhibition and stimulation of electrical activity in the HH models under study. This framework relies only on the bifurcation analysis of equilibrium points of the system, thereby avoiding the computational burden required by the study limit of cycles. We briefly took into account also the possibility of performing the ionic actuation locally at the cell membrane. To this aim, we considered a two-compartment model to lump the actuated and non-actuated patches of the membrane, along with their electrical coupling. We found that the portion of actuated membrane *ρ* is a critical parameter since a value of Δ*V*_K_ that elicits a certain regime in the single-compartment case must be increased by a factor comparable to *ρ*^−1^ to induce the same behavior in the two-compartment case. The strength of electric coupling *g_c_* modulates such factor, and may introduce novel steady regimes in the system.

A large body of literature points out K^+^ as a modulator of electrical activity in neurons, both in experiments [7–10] and in computational studies [11–18]. These works are mainly concerned with the regulation of ionic concentrations and pathological conditions that result in their impairment, such as seizures and spreading depression. Since such phenomena occur at time scales of seconds or even minutes, synaptic currents are not typically considered as bifurcation parameters. In a neuromodulation scenario, such analysis is instead pivotal since synapses act as uncontrolled input of the system and the time scale of the intrinsic electrical activity of neurons is a major concern. In [39], a bifurcation diagram similar to the ones in Fig. 5 is reported. However, a dual perspective on the problem is assumed. In fact, in that reference, the current is the main driver of neural activity, while the perturbation of [K^+^] is a consequence of prolonged firing. The paper thus reports a switching between class I and class II excitability, which is verified *in-vitro* and is consistent with our findings. As a complement to the previous works, we show that the switchings of the class of excitability depend on the type of neuron considered. Moreover, we report that also switching to class III is possible. In the latter case, eliciting tonic spiking is not possible if ionic actuation is assumed as the sole input of the system. Our analysis also offers a comparative computational study of the differential impact of [K^+^] on the electrical activity of neurons of different classes. Building upon these findings, one may expect that neurons *in-vitro* or in culture may show or not transition to a spiking behavior, or require different extents of ion release to enter into block. Such speculations might motivate new directions for experimental investigations.

There are a number of caveats to consider here. First, we described the action of the ionic actuator solely as a perturbation of extracellular K^+^ concentration. However, the dynamics of ions involve electrodiffusive processes that may influence also the electric field in the extracellular microenvironment, thereby introducing further paths along which ionic actuation may affect the cell’s electrical activity. If such effects are not negligible, a more accurate description of the ionic actuator must be included in the analysis. Second, we considered reduced HH models for the electrical activity of neurons. The single-compartment models have been fitted to reproduce the electrophysiology of actual neural cells and, therefore, might provide a reasonable understanding of the effects of an ionic actuator that is able to perturb all the surrounding milieu of the cell. However, the two-compartment model is rather artificial and was introduced to suggest possible key ingredients affecting a local ionic actuation of the cell, rather than to reproduce actual electrophysiological properties in this setting. Indeed, the change in *V*_K_ between the two compartments does not account for any spatial gradient that may arise from the diffusion of ions, and the electric coupling between the actuated and non-actuated regions is mimicked with a phenomenological parameter *g_c_*. Therefore, to draw a more realistic insight into such scenario, it is necessary to resort to models that include more explicitly the spatial nature of the phenomena. This will enable also the identification of “hot zones” where the modulation of the neuron is facilitated, such as the well-known initial segment for spike initialization [46].

We focused on the ionic neuromodulation of a neuron. A natural generalization of such application may be the control of a neuronal network. However, this introduces several complications since neurons form networks where both excitatory and inhibitory neurons are present. Indeed, in a single-cell scenario to silence (respectively, trigger) electrical activity, it is sufficient to drive the system to either rest or block (respectively, spiking). Conversely, in a network scenario, the steering of a neuron in a rest or block state may reduce or increase the overall electrical activity in the network, depending on whether it is part of the excitatory or inhibitory class. The converse applies to a neuron that is driven to a spiking regime. It is not a surprise that these interactions underlie several epileptic events [47–51], where the perturbation of the electrical activity of neurons is induced by pathological conditions rather than neuromodulatory devices. The scenario is further complicated by the previously mentioned differential influence of [K^+^]_*o*_ on the electrical activity of different types of neurons. The ionic neuromodulation of an ensemble of neurons is therefore a more involved topic, which at the same time permits to envisage a much richer ensemble of control patterns to steer such networks. It is thus clear that to approach its study one cannot ignore the network interactions, as well as the intrinsic peculiarities of each cell.

In conclusion, we provided a theoretical insight into the employment of the extracellular concentration of K^+^ as a novel neuromodulation modality. Leveraging bifurcation analysis, we highlighted the possibilities and limits offered by such modality in shaping the electrical activity of neurons. We showed through simulations how these findings may inform control strategies aiming to either stimulate or inhibit electrical activity. Our results complement the advances in the technology of ionic actuators and foster the design of novel ionic-enabled neural interfaces.

## Matlab code

The code is available publicly at github.com/claudioverardo/neuro-ionact and it is implemented in Matlab [42]. The simulations were made with the built-in `odel5s` solver. The bifurcation analyses of equilibrium points were made as described in the section Methods, exploiting the generalizations presented in Appendix B. The bifurcation analyses of limit cycles were made with the open-source toolbox MatCont [43].

## Appendix A: Hodgkin-Huxley-type models

In our investigations, we considered some HH-type models from the literature as case studies. These are listed in Table 1. Their membrane parameters and the kinetics of their gating variables are reported in Tables 3 and 4, respectively. These models comprise two voltage-gated channels for potassium and sodium, denoted as K and Na, responsible for the generation of action potentials. The models rat-pospischil08-FSin and rat-pospischil08-RSexch also include a muscarinic potassium channel, denoted as K*_m_*, responsible for spike adaptation. The models rat-wei14 and rat-cressman09 also include ion-specific leakage channels for potassium and sodium, denoted as K_*L*_ and Na_*L*_. For the sake of simplicity, in the mathematical derivations of the main body of the paper, we considered only the channels K and Na. However, such derivations readily generalize to include further channels, as shown in the Appendix B.

**Table 3.** Parameters of the HH-type models used as case studies. In all the models, the membrane capacitance is *C_m_* = 1 μF/cm^2^. In squid-hh52, temperature correction factors are applied to both the kinetics of the gating variables 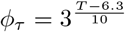, and to the conductances 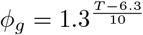 according to [52]. In rat-wang96, the original work [40] does not state the temperature *T* explicitly, but reports experimental data recorded at 35 – 37 °C.

| ↓ Models Parameters → | $V_r$ | $V_{K,0}$ | $V_{Na}$ | $V_L$ | $\bar{g}_K$ | $\bar{g}_{K_m}$ | $\bar{g}_{Na}$ | $g_{K,L}$ | $g_{Na,L}$ | $g_L$ | $T$ | $\phi$ |
| --- | --- | --- | --- | --- | --- | --- | --- | --- | --- | --- | --- | --- |
| squid-hh52 [34] | -60 | -76 | 55 | -44.5 | $36 \phi_g$ | - | $120 \phi_g$ | - | - | $0.3 \phi_g$ | 20 | $\phi_\tau$ |
| rat-wei14 [14] | -66.8 | -94.7 | 55.4 | -81.9 | 25 | - | 30 | 0.05 | 0.0175 | 0.05 | 36 | 1 |
| rat-cressman09 [11] | -67.0 | -94.7 | 55.4 | -81.9 | 40 | - | 100 | 0.05 | 0.0175 | 0.05 | 36 | 3 |
| rat-wang96 [40] | -64 | -90 | 55 | -65 | 9 | - | 35 | - | - | 0.1 | 37 | 5 |
| rat-pospischil08-FSinh [41] | -71.4 | -90 | 50 | -70.4 | 3.9 | 0.079 | 58 | - | - | 0.038 | 36 | 1 |
| rat-pospischil08-RSexc [41] | -71.9 | -90 | 50 | -70.3 | 6 | 0.075 | 56 | - | - | 0.021 | 36 | 1 |
| Units → | mV |  |  |  | mS/cm<sup>2</sup> |  |  |  |  |  | °C | - |

**Table 4.**
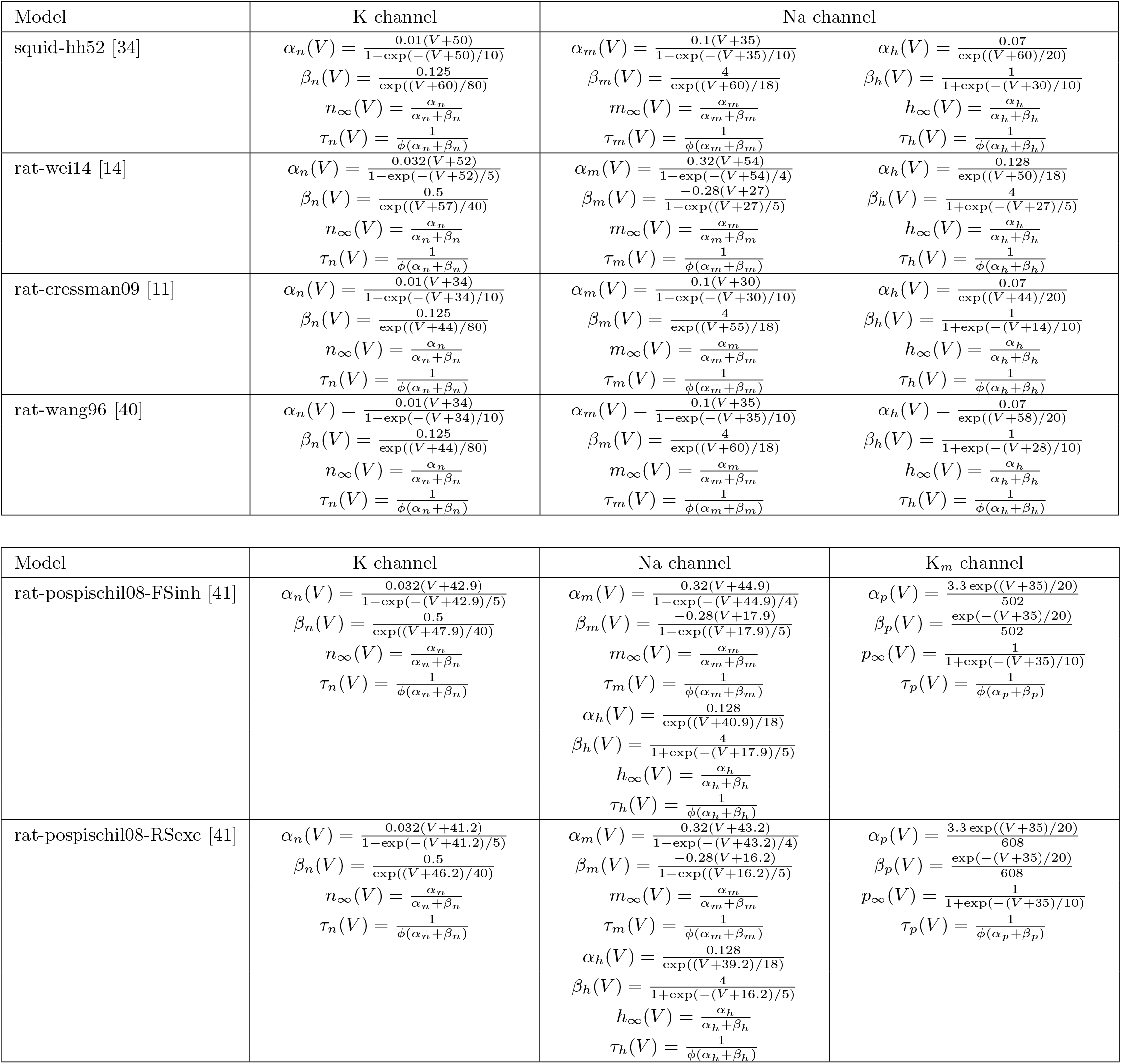
Kinetics of the HH-type models used as case studies. The models squid-hh52, rat-wei14, rat-cressman09, rat-wang96 (top) only contain a K channel and a Na channel responsible for the generation of action potentials. In the models rat-pospischil08-FSinh and rat-pospischil08-RSexc (bottom), there is an additional muscarinic K channel (K*_m_*) responsible for spike adaptation. The conductance of the K channel is 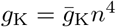, while the conductance of the Na channel is 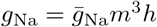. When present, the conductance of the muscarinic K channel is 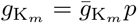. The values for the density of channels 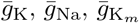, and the temperature coefficient *ϕ* are reported in Table 3. In the main body of this paper, for the sake of simplicity, only K and Na channels are assumed in the mathematical derivations. The authors refer the reader to Appendix B for a generalization of such derivations to other channels such as K*_m_*.

## Appendix B: generalization to arbitrary ion channels

The derivations presented in the main body of this paper readily generalize to HH models with more channels than action potential-generating K and Na channels. Let us consider a model with *N_ch_* voltage-gated channels and *N_sp_* ionic species involved in the transmembrane fluxes. *g_i_* denotes the conductance of the *i*-th voltage-gated channel. X*_j_* denotes the *j*-th ionic species (e.g., K^+^, Na^+^, Ca^2+^, Cl^−^), *V*_X_*j*__. is the Nernst potential of X_*j*_, and *g*_X_*j*__,*L* is the X_*j*_-specific leakage conductance. We point out that more channels may share the same Nernst potential (e.g., the K and K*_m_* channels in Table 4). In the case of non-specific channels involving cations, the contribute of K^+^ should be properly split to become sensible to ionic actuation. We define:

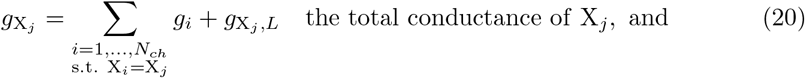

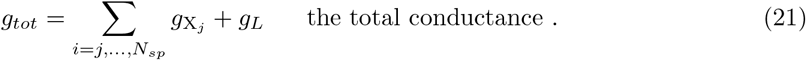

The conductance of the *i*-th voltage-gated channel is assumed in the form 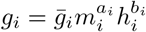, where *m_i_* is the activating gating variable and *h_i_* the inactivating gating variable. When such quantities are evaluated at steady-state, we adopt the subscript “∞”. We neglect possible dependencies other than the membrane potential in the kinetics of the gating variable and in the channel conductances, e.g., concentrations of neurotransmitters or calcium. Therefore, the equations modeling the dynamics of the system are

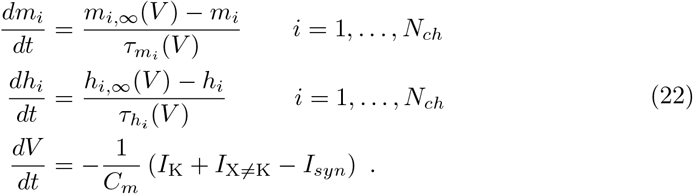

Here, *I*_K_ is the current associated with K^+^-selective channels, while *I*_X≠K_ is the current associated with the remaining channels. Namely,

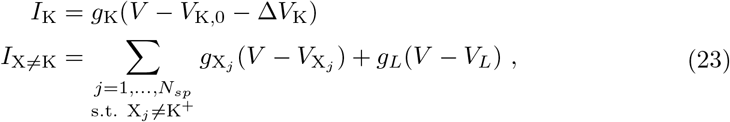

where *g*_K_ is defined according to Eq. (20) with X_*j*_ = K^+^. The equilibrium points of the system (22) are in the form (…, *m*_*i*,∞_(*V_s_*), *h*_*i*,∞_(*V_s_*), …, *V_s_*), with *V_s_* sets by

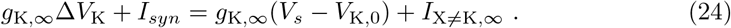

Therefore, maps as Eqs. (7) and (8) still hold, with 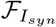 and 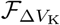 now defined as

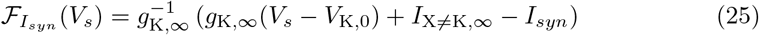

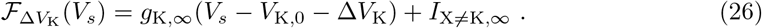

The Jacobian takes the form of a block matrix. Namely,

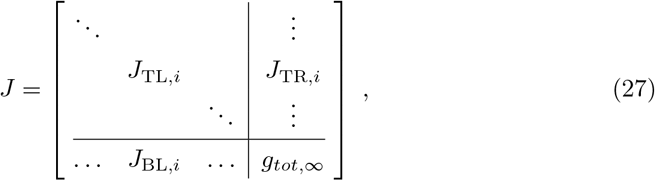

where, for *i* = 1, …, *N_ch_*, the blocks *J*_TL,*i*_, *J*_TR,*i*_, *J*_BL,*i*_ have dimensions 2 × 2, 2 × 1, 1 × 2, respectively, and are in the form

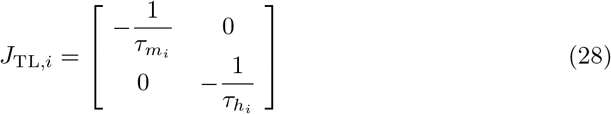

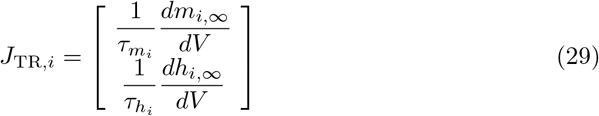

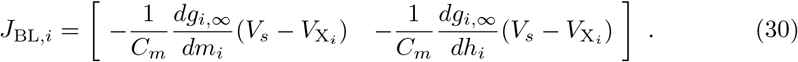

Studying the eigenvalues of *J* is possible to find the bifurcations *V_th_* and *V_block_* as shown in the main body of the paper. The maximum dimension of *J* is 2*N_ch_* × 2*N_ch_*. However, if a gating variable is not present (*a_i_* = 0 or *b_i_*=0), the corresponding entries in *J* must be removed, and the number of eigenvalues is reduced accordingly.

The first-order transfer functions *A_I_* and *A_V_* can be generalized by linearizing Eq. (24). The relation *A*_K_ = *g*_K,∞_*A_I_* still holds, but now *A_I_* reads

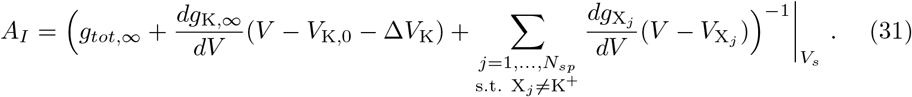

## Acknowledgments

We thank Federico Leva and Prof. Luca Selmi from University of Modena and Reggio Emilia for the support and the fruitful discussions. We acknowledge funding from the European Unions Horizon 2020 research and innovation programme under grant agreement No 862882 (IN-FET project) via the IUNET Consortium.

